# Frequency-specific neuromodulation of local and distant connectivity in aging & episodic memory function

**DOI:** 10.1101/061267

**Authors:** Simon W. Davis, Bruce Luber, David L. K. Murphy, Sarah H. Lisanby, Roberto Cabeza

**Author notes:** Corresponding author: Simon W Davis, Department of Neurology, Box 2900, DUMC, Duke University, Durham, NC 27708.

## Abstract

A growing literature has focused on the brain’s ability to augment processing in local regions by recruiting distant communities of neurons in response to neural decline or insult. In particular, both younger and older adult populations recruit bilateral prefrontal cortex (PFC) as a means of compensating for increasing neural effort to maintain successful cognitive function. However, it remains unclear how local changes in neural activity affect the recruitment of this adaptive mechanism. To address this problem, we combined graph theoretical measures from functional MRI (fMRI) with diffusion weighted imaging (DWI) and repetitive transcranial magnetic stimulation (rTMS) in order to resolve a central hypothesis: *how do aged brains flexibly adapt to local changes in cortical activity?* Specifically, we applied neuromodulation to increase or decrease local activity in a cortical region supporting successful memory encoding (left dorsolateral prefrontal cortex or DLPFC) using 5Hz or 1Hz rTMS, respectively. We then assessed a region’s local *within-module degree* (WMD), or the distributed *between-module degree* (BMD) between distant cortical communities. We predicted that (1) local stimulation-related deficits may be counteracted by boosting BMD between bilateral PFC, and that this effect should be (2) *positively correlated* with structural connectivity. Both predictions were confirmed; 5Hz rTMS increased local success-related activity and local increases in PFC connectivity, while 1Hz rTMS decreases local activity and triggered a more distributed pattern of bilateral PFC connectivity to compensate for this local inhibitory effect. These results provide an integrated, causal explanation for the network interactions associated with successful memory encoding in older adults.

## Introduction

Over the past few decades, transcranial magnetic stimulation (TMS) has developed into a powerful tool to causally establish brain-behavior relationships. Traditionally, inferences on the consequence of TMS have been limited by the fact that it can only directly target superficial cortex. Studies combining TMS and fMRI or EEG have demonstrated that TMS can result in measurable effects at cortical sites both local and distant to the site of stimulation (Bestmann et al., 2004; Wang et al., 2014), suggesting that the effects of TMS should be considered within the context of a more interconnected global network. While cognitive functions like episodic memory encoding are now known to be supported by a largely distributed network of interacting cortical regions, an understanding of how regional and frequency-specific manipulations of TMS stimulation affect these distributed network is lacking. The goal of the present study is to investigate the frequency-specific responses of memory-related cortical networks in response to local neurostimulation. Furthermore, it is unclear how differences in stimulation protocols map onto these global network dynamics, or how cognitive states can be selectively targeted using dynamic spatiotemporal signals distributed over large-scale networks of the brain.

We focused on the well-known phenomenon that older adults tend to over-recruit an more distributed pattern of bilateral prefrontal cortex (PFC) regions to those most active in younger adults. Age-related increases in contralateral recruitment have been observed many times across a variety of studies using different stimuli and cognitive tasks, including episodic encoding (Persson et al., 2006), semantic retrieval (Davis et al., 2012), attention (Huang et al., 2012), motor coordination (Heuninckx et al., 2008), and working memory (Reuter-Lorenz et al., 2000). This effect follows the more general observation of age-related reductions in core task-network regions coupled with an increase in secondary task-network areas (Dennis and Cabeza, 2008; Nyberg et al., 2010), but may also be a specific example of adaptive efficiency in the aging brain. While the prevalence of this pattern in aging studies is widespread, the evidence that this effect benefits performance is mixed, and contralateral recruitment may only be effective if the additional cortical processors brought to bear on the task can play a complementary role in task performance (Colcombe et al., 2005). Such a model of the aging brain proposes that over the years, cortical messages are increasingly dominated by top-down predictions, and less by sensory inputs (Moran et al., 2014). This is consistent with the age-related shift to PFC activation, and bilateral PFC activity is a likely consequence of this adaptive shift, given that every PFC region is connected to its contralateral homologue by at least one synapse (Aboitiz et al., 1992; Aboitiz and Montiel, 2003). Nonetheless, while correlational studies have observed that PFC regions over-recruited by OAs often show stronger long-range functionally connectivity outside of local cortical communities (Davis et al., 2012; Geerligs et al., 2015) and have been associated with successful cognitive performance in this group (Dennis et al., 2008; Spaniol and Grady, 2012), the adaptive dynamics of this effect are unclear. We and others have suggested that OAs compensate for local deficits by expanding the task-related network via long-range connectivity to bilateral cortical systems (Antonenko et al., 2012; Davis et al., 2012; Burzynska et al., 2013). The bilateral PFC system observed in older adults therefore serves as useful model with which to explore the adaptive mechanisms of local and global communication in response to TMS. Our approach is therefore to unite network theory, brain stimulation, and diffusion tractography, in order determine how the aging brain adapts to local inhibition or excitation of cortical reactivity.

Noninvasive brain stimulation technologies offer a unique opportunity to probe the functional and structural reorganization to resolve the functional relevance of network structure. Repetitive transcranial magnetic stimulation (rTMS) at higher frequencies (≥ 5Hz) has been found in induce local *increases* in BOLD activity, during a host of different cognitive operations including motor planning (Schneider et al., 2010), working memory (Esslinger et al., 2014), and episodic memory (Vidal-Pineiro et al., 2014), *boosting* ongoing activity associated with successful performance. Conversely, 1Hz rTMS has been shown to reliably *depress* local hemodynamic activity (de Vries et al., 2012; Plow et al., 2014) and impairs cognitive performance associated with the local region (Binney et al., 2010). While the use of rTMS as a tool for behavioral neuroenhancement is undoubtedly informed by understanding these local effects, it remains unclear how local changes in neural activity affect large-scale system dynamics. If the age-related increase in distal connectivity patterns reflects a compensatory phenomenon, we would expect this 1Hz TMS to induce an increase in distal connectivity associated with successful memory.

Here we propose and test a new pair of hypotheses based on the graph measures of modularity (Chang et al., 2012) and degree. Network modularity is defined by the existence of distinct groups of nodes which connect more intimately with each other than with other nodes in the network. A modular decomposition allows one to then consider the behavior of the strength of connections within a *local* or more *global* community of regions. Once the modular *network architecture* has been established by a reliable means, the extent of local or global *network activity* can be assessed quantitatively using *within-module degree* (WMD) and *between-module degree* (BMD). WMD represents the sum of local connectivity at a node with all of the nodes in its local cortical module, whereas BMD describes the sum of connectivity between a local node and nodes within other *distinct* modules. A more well-known measure of extra-module connectivity, participation coefficient (PC), severely reduces the specificity of distant connectivity by summing degree values across all non-local communities, making it difficult to assess what distant communities are being recruited. Using these ideas, we examine the widespread activation pattern changes in long-range functional connectivity displayed in response to local modulation by rTMS, in a group of healthy older adults. In the current study, we use TMS to test two principle predictions. First, we directly manipulate the hypothetical cause of the compensatory change, i.e., the operation of a task-related brain region associated with successful memory encoding. Using rTMS we either impair (1Hz) or enhance (5Hz) the function of core region for verbal episodic memory encoding: left PFC. We predict that *impairing left PFC function will increase BMD between distant PFC regions, whereas enhancing its function will increase local WMD with the stimulation site (Prediction 1)*. Second, given that functional connectivity depends on structural connectivity (Gong et al., 2009), adaptive changes in functional connectivity after neuromodulation should be predicted by the availability of white-matter pathways between task-related nodes in a network that allow such adaptation to occur. Therefore, *we predict that while WMD will be correlated with direct structural connections with the stimulation site, BMD will be related to indirect structural connectivity* (*Prediction 2*).

To test these predictions, we establish the map of local and distant topological communities by their static, structural connectivity patterns, as given by diffusion tractography. We delivered both concurrent TMS/fMRI, as well as 1Hz and 5Hz rTMS immediately before scanning while encoding sentences, in order selectively inhibit (1Hz) or excite (5Hz) the local area of activity related to successful memory encoding. The first prediction was tested by comparing the two rTMS conditions on WMD and BMD; the second prediction was tested by correlating WMD and BMD with structural distance measures. The confirmation of these three predictions would directly support our account of widespread activity in OAs.

## Materials & Methods

### Participants

Fifteen healthy older adults were recruited for this study (all native English speakers; 8 females; age mean +/- SD, 67.2 +/- 4.4 years; range 61-74 years); one subject was subsequently removed from the analysis due to tolerability during rTMS. Each older adult was screened for exclusion criteria for TMS (history of seizure, brain/head injuries) as well as psychiatric condition (MINI International Neuropsychiatric Interview, English Version 5.0.0 DSM-IV, Sheehan et al., 2006). None of the older participants reported subjective memory complaints in everyday life or had MMSE score below 27 (mean +/- SD = 29.1 +/- 0.8).

### Stimuli & Procedure

The source memory task was comprised of a set of 360 sentences, each of which included a concrete subject and direct object. In each sentence, both subject and direct object were capitalized to indicate to the subject which specific nouns were to be remembered (*“A SURFBOARD was on top of the TRUCK.”*). Associative strength between nouns in a sentence (as determined by the USF word association norms (Nelson et al., 2004)), were normally distributed, and both the imageability, frequency, and total length of each set of sentences was counterbalanced across all sentences used therein.

One run of the source task comprised an encoding block of 90 sentences presented visually for 3 seconds, and a subsequent retrieval test comprising 68 word pairs from the same previously studied sentence, and 22 new pairs composed of two words recombined from two different sentences. Participants were asked to judge whether each word pair was intact or recombined and indicate how confident they were in their decision on a 4-point rating scale. Given the low proportion of low-confidence responses in the current data, we collapsed low and high-confidence responses, and excluded encoding trials in which subjects failed to indicate a response at retrieval.

### Image Acquisition

An outline of all data acquisition events is depicted in Figure 1. Scanning was divided between two days, 1-4 days apart, with the first day comprised of a functional memory-success localizer for scanning and stimulation on Day 2. All procedures were completed on a GE MR 750 3-Tesla scanner (General Electric 3.0 tesla Signa Excite HD short-bore scanner, equipped with an 8- channel head coil). Coplanar functional images were acquired with an 8-channel head coil using an inverse spiral sequence with the following imaging parameters: flip angle = 77°, TR = 2000ms, TE = 31ms, FOV = 24.0 mm^2^, and a slice thickness of 3.8mm, for 37 slices. The diffusion-weighted imaging (DWI) dataset was based on a single-shot EPI sequence (TR = 1700 ms, 50 contiguous slices of 2.0 mm thickness, FOV = 256 × 256 mm2, matrix size 128 × 128, voxel size 2 × 2 × 2 mm, b-value = 1000 s/mm2, 25 diffusion-sensitizing directions, total scan time ∼5 min). The anatomical MRI was acquired using a 3D T1-weighted echo-planar sequence (256 × 256 matrix, TR = 12 ms, TE = 5 ms, FOV = 24 cm, 68 slices, 1.9 mm slice thickness). Scanner noise was reduced with earplugs and head motion was minimized with foam pads. Behavioral responses were recorded with a four-key fiber optic response box (Resonance Technology), and when necessary, vision was corrected using MRI-compatible lenses that matched the distance prescription used by the participant.

**Figure 1.**
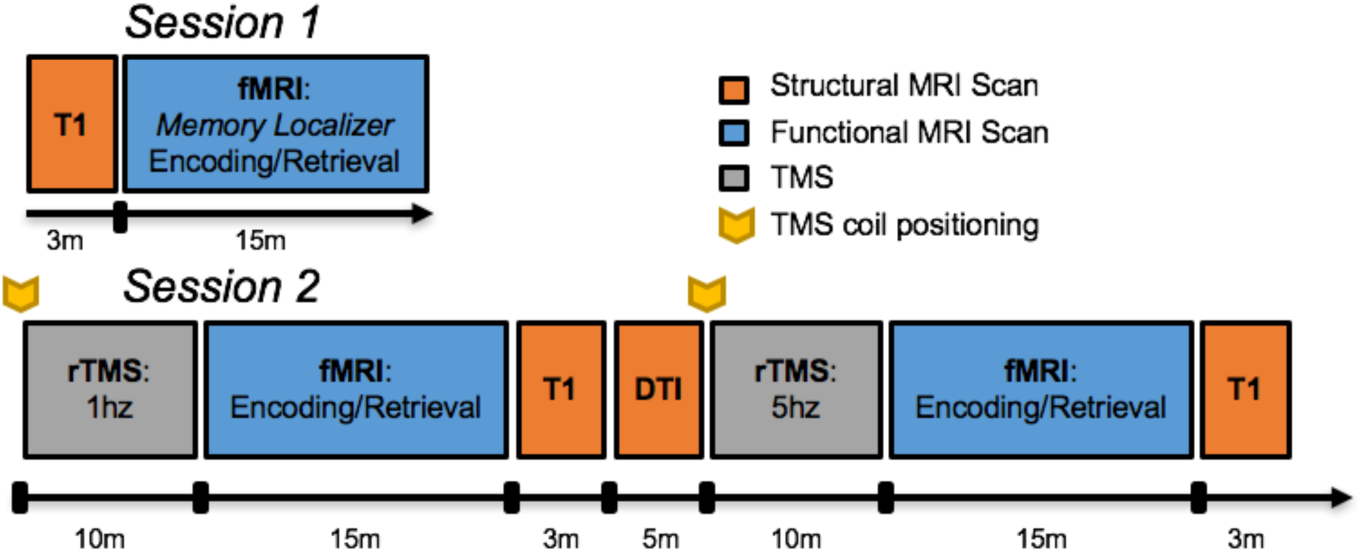
Data acquisition schedule.

### TMS Procedure

Prior to the TMS-fMRI session, the scalp location for TMS coil position over the left middle frontal gyrus (MFG) location had been found by using an infrared neuronavigation system (Brainsight: Rogue Research, Montreal, Canada). Specifically, the point of greatest activation in the left MFG in the fMRI memory contrast (i.e., encoding trials which were subsequently remembered versus forgotten) from the first day of scanning was chosen from the fMRI overlay on the subject’s structural MRI, both of which had been uploaded into BrainSight. After co-registration of the subject’s head with his MRI, the MFG location was marked on a tight fitting acrylic swim cap that stayed on the subject’s head until TMS-fMRI interleaving was completed on the same day. At that time, subjects were acclimated to the sort of TMS pulses to be delivered later in the scanner with a series of single pulses at the target site, as well as a short burst of 5Hz stimulation. The motor threshold (MT) for each subject was determined using a MagVenture R30M device located outside the scanner room, part of an MRI compatible TMS system which included a non-ferrous figure-8 coil with 12m long cable and artifact reducing counter-current charging system (MagVenture, Farum, Denmark). MTs were determined using electromyography of the right first dorsal interosseous (FDI) muscle and defined as the lowest setting of TMS device intensity at which ≥ 5 out of 10 motor evoked potentials of at least 50µV peak to peak amplitude could be elicited.

Before each functional scan, two 10-minute trains of either 1Hz or 5Hz stimulation were delivered at 120% MT immediately prior to a fMRI acquisition. The position of the TMS coil was reset to the same target site before the beginning of each rTMS session, and monitored continuously while the subject lay supine in the bed of the MR scanner. 1Hz rTMS was delivered in a continuous train of 10 minutes, while 5Hz rTMS was delivered in intermittent 6 sec trains with a 24 sec inter-train interval, also for 10 minutes. Dosage was equivalent between 1Hz and 5Hz rTMS conditions (600 total pulses), and the order of stimulation frequency was counterbalanced across subjects. Immediately after the 10 minutes of TMS, subjects were positioned in the scanner, and performed the encoding and subsequent retrieval portion of the sentence task while fMRI was acquired. The amount of time elapsed between the end of the rTMS train to the beginning of the functional scan was 9.4 minutes (SD = 1.7 minutes).

### Data Analyses

The general analytical pipeline is depicted in Figure 2. Our general approach is to collect both functional and structural imaging data, and create adjacency matrices comprising psychophysiological interaction (PPI)-based functional correlations associated with the task (fMRI data) or structural connections based on tractography streamline counts (DWI data). Modularity was based solely on structural connectivity, and regions within a high-dimensional atlas (n = 411 regions) were partitioned using conditional expected models (Chang et al., 2012). In order to compare functional and structural networks we use graph measures to find the distance traversed by the lowest weighted path in a structural network, which is constructed by connecting each cortical region with a weighted edge. by which we are able to demonstrate the value of both local (direct) and long-distance (indirect) paths in promoting success-related connectivity.

**Figure 2.**
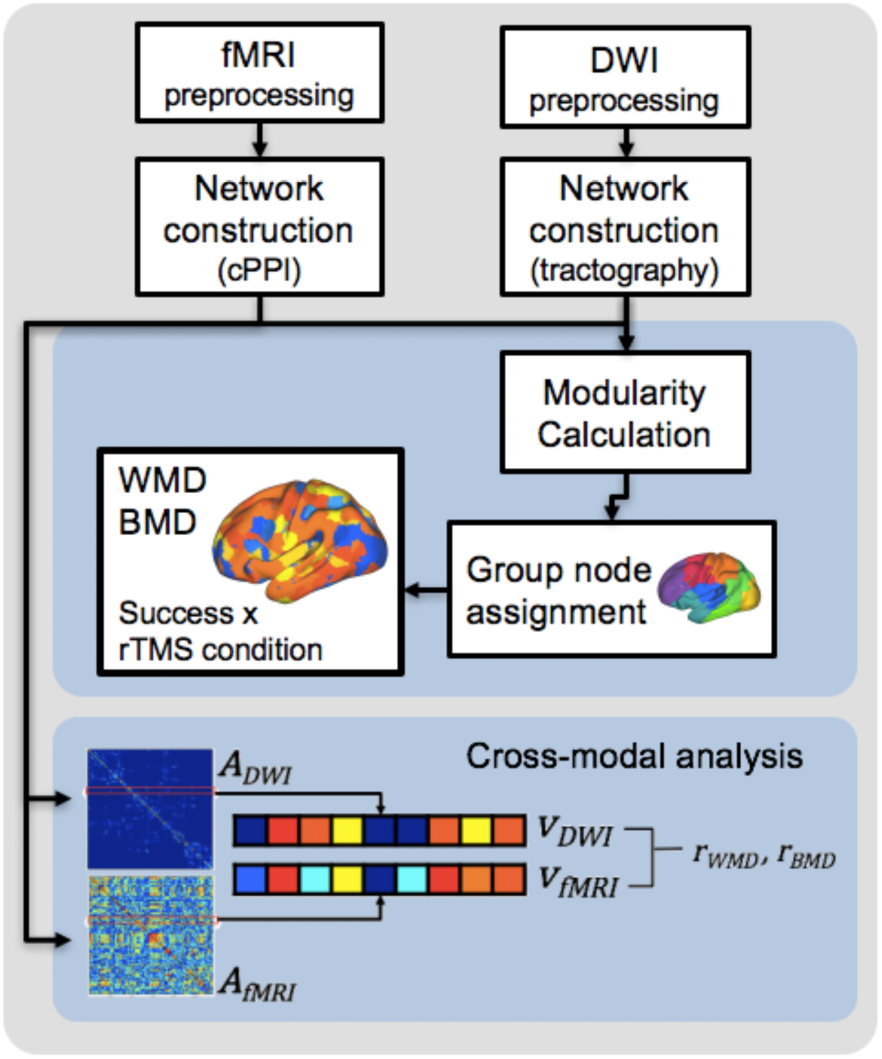
Analytical pipeline. Schematic depicts the general analytic pipeline used in the current study, both functional and structural imaging data were processed according to standard preprocessing heuristics (see Material & Methods), and adjacency matrices comprising functional correlation associated with the task (fMRI data) or structural connections based on streamline counts (DWI data) were evaluated. Then, both structural and functional matrices are submitted to a modular decomposition. Next, the modal decomposition across all subjects is used to describe the most common modular breakdown (in the baseline Encoding Success (DM) condition). Within-module degree (WMD) and between-module degree (BMD) are then computed for each region. Lastly, cross-modal analyses are completed by calculating the Pearson’s product moment correlation *r*, based on vectors representing connectivity information from either within- or between-module connections.

### Structural and functional preprocessing

Structural diffusion-weighted images were preprocessed using standard DWI preprocessing, including brain extraction, correction for eddy-current distortion and simple head motion, and correction of the b-matrix for any rigid-body coregistration, using the *bet, eddy,* and *FNIRT* functions from FSL (FMRIB, Oxford, UK), respectively. Functional images were preprocessed using image processing tools, including FLIRT and FEAT also from FSL, in a publically available analysis pipeline developed by the Duke Brain Imaging and Analysis Center (https://wiki.biac.duke.edu/biac:analysis:resting_pipeline). Images were corrected for slice acquisition timing, motion, and linear trend; motion correction was performed using FSL’s MCFLIRT, and 6 motion parameters estimated from the step were they regressed out of each functional voxel using standard linear regression. Images were then temporally smoothed with a high-pass filter using a 190s cutoff, and normalized to the Montreal Neurological Institute (MNI) stereotaxic space. White matter and CSF signals were also removed from the data, using WM/CSF masks generated by FAST and regressed from the functional data using the same method as the motion parameters. Spatial filtering with a Gaussian kernel of full-width half-maximum (FWHM) of 6mm was applied. Voxel time-series analysis was carried out using general linear modeling (GLM); fixed effects models were carried out to examine the effects of both memory success and stimulation frequency in the baseline and post-rTMS data. We modeled encoding trials that were either subsequently remembered or subsequently forgotten, in order to examine standard subsequent memory effects (SMEs) in the baseline, post-1Hz rTMS, and post-5Hz rTMS runs.

### Construction of connectivity matrices

Before either structural or functional matrices were constructed, we first sought to establish a consistent parcellation scheme across all subjects and all modalities (DWI, fMRI) that reflects an accurate summary of full connectome effects (Bellec et al., 2015). Subjects’ T1-weighted images were segmented using the SPM12 (<u>www.fil.ion.ucl.ac.uk/spm/software/spm12/</u>), yielding a grey matter (GM) and white matter (WM) mask in the T1 native space for each subject. The entire GM was then parcellated into 411 regions of interest (ROIs), each representing a network node by using a subparcellated version of the Harvard-Oxford Atlas, (Tzourio-Mazoyer et al., 2002), defined originally in MNI space. The T1-weighted image was then nonlinearly normalized to the ICBM152 template in MNI space using fMRIB’s Non-linear Image Registration Tool (FNIRT, FSL, <u>www.fmrib.ox.ac.uk/fsl/</u>). The inverse transformations were applied to the HOA atlas in the MNI space, resulting in native-T1-space GM parcellations for each subject. Then, T1-weigted images were coregistered to native diffusion space using the subjects’ unweighted diffusion image as a target; this transformation matrix was then applied to the GM parcellations above, using FSL’s FLIRT linear registration tool, resulting in a native-diffusion-space parcellation for each subject.

For structural connection matrices, network edges were defined by the number of tractography streamlines between any two ROIs. We used Dipy (Garyfallidis et al., 2014) to fit the data with the constant solid angle (CSA) model with fourth-order spherical harmonics, and generating generalized FA (GFA) maps from estimated orientation distribution functions. Using a deterministic tracking algorithm (EuDX; Garyfallidis et al., 2012) with standard tracking parameters (step size: 0.5mm, turning angle 45**°**, alpha = 0.05), within native-space. Whole-brain streamlines (∼30,000 per subject) were saved for subsequent parcellation into structural connectivity matrices. As the current analysis Lastly, we corrected for spurious node pairs using a correction method described previously (Gong et al., 2009), in which a nonparametric sign-test was applied by taking each individual as a sample, with the null hypothesis being that there is no existing connection, and a node pair surviving a corrected p < 0.05 was deemed to have a connection; all node pairs below this threshold were excluded from subsequent analyses.

Functional connection matrices representing task-related connection strengths were estimated using a correlational psychophysical interaction (cPPI) analysis (Fornito et al., 2012). Briefly, the model relies on the calculation of a PPI regressor for each region, based on the product of that region’s timecourse and a task regressor of interest, in order to generate a term reflecting the psychophysical interaction between the seed region’s activity and the specified experimental manipulation. In the current study the task regressors based on the convolved task regressors from the univariate model described above were used as the psychological regressor, which coded subsequently remembered and subsequently forgotten word pairs with positive and negative weights, respectively, of equal value. This psychological regressor was multiplied with two network timecourses for region *i* and *j*. We then computed the partial correlation 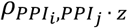, removing the variance *z* associated with the psychological regressor, the timecourses for regions *i* and *j*, and constituent noise regressors. We accounted for the potential effects of head motion and other confounds by assessing the 6 motion parameters and including these parameters in our partial correlation between regions.

### Graph Theory Metrics

The network modeling approach used in the current manuscript focuses on two key mechanistic concepts: modularity and degree. Degree is a basic graph measure describing the sum of edges at a node *i* in weighted networks, while modularity is a measure originally derived for quantifying the quality of a partition (Newman and Girvan, 2004), but more recently has become a key graph measure for quantifying the relationship both within and between subnetworks in a whole-brain graph. We used two well-established connectivity metrics, *modularity* and *within-module degree* (WMD), and one novel metric, *between-module degree* (BMD). Additionally, we aimed to validate BMD with an established measure of extra-module connectivity, namely participation coefficient (PC). Modularity (both Q and nodal assignments), WMD, BMD, and PC were computed with a combination of tools from the Brain Connectivity Toolbox (Rubinov and Sporns, 2010) and custom MATLAB scripts. We discuss these measures in more detail below.

#### Modularity

We chose to use subjects’ structural connectivity (DWI tractography) to define modularity across subjects; While the use of ‘canonical’ functional partitions based on resting-state data (e.g., Power et al., 2011) is growing in popularity, owing both to their internal reliability and ease of data collection, it is still unclear how appropriate such parcellations may be for task-based connectivity (for review of this issue, see Davis et al., 2016). In the current study, resting state correlations may, and modularity based on task-related connectivity is problematic, given the uncertainty in how neuromodulation may affect these measures. Thus, our more conservative approach was to use structural connectivity (DWI tractography) to define community membership (i.e., modularity).

Fundamental to the definition of modularity is the computation of a proper null network, with the same number of nodes and node degrees but otherwise no underlying structure. Standard modularity analysis relies on defining a null distribution of possible connections from a randomization of the rows of the input matrix (Newman, 2006). However, even random networks can exhibit high modularity because of incidental concentrations of edges, and this method of computing the null is also problematic for our analysis because it ignores negative connection weights (i.e., anti-correlated regions) and implicitly assumes self-loops (connections from nodes to themselves), which are meaningless in the functional networks considered here. Furthermore, it is widely observed (but rarely reported) that modularity partitions based on randomized null distributions suffer from inconsistency in partition assignments over repeated executions of the same algorithm, resulting in different partitions *with each repetition of the modularity algorithm*. To reach some standard of replicability, researchers have instead come to rely on permuting the algorithm for the maximum *Q*, or modularity quotient. We have therefore sought to employ a more robust modularity algorithm (Chang et al., 2014) that relies on a transformed Tracy-Widom distribution in order to more adequately model the null distribution in a modularity computation. Furthermore, while it is possible to estimate modularity within individuals (e.g., Simpson et al., 2012), modularity based on an average are naturally consistent across subjects, and widely used (Hagmann et al., 2008; Lim et al., 2015). We therefore estimated a group DTI connectivity matrix based on the mean of streamline counts across all subjects, for each element of the 411 × 411 matrix, and the computed modularity based on this average matrix. Such a consistent parcellation facilitates the interpretation of the local versus distant connectivity changes induced by different frequencies of rTMS observed below.

#### Within-Module and Between-Module Degree

The sum or connections (or edges) between a node and other nodes in the network defines the node’s degree. We partitioned these degree values either within all nodes within a particular module (WMD), or between a node and all other cortical modules (BMD). As such, WMD and BMD help to characterize the local and distributed connectivity patterns during successful memory formation. Note that *distributed*, in this context, is defined by the topological properties of a connection traversing the modules calculated above, and not necessarily the Euclidean distance between nodes.

Following previous studies focusing on degree (Guimera and Nunes Amaral, 2005), we evaluated WMD and BMD degree distributions between encoding trials which were subsequently *remembered* or *forgotten*, in both post-rTMS conditions (1Hz and 5Hz rTMS). Critically, we considered these changes in degree distribution at each node either with respect to its surrounding module (within-module degree or WMD), or as a function of a node’s relationship with more distant cortical modules (between-module degree or BMD). Both WMD and BMD use the same underlying function to estimate degree:

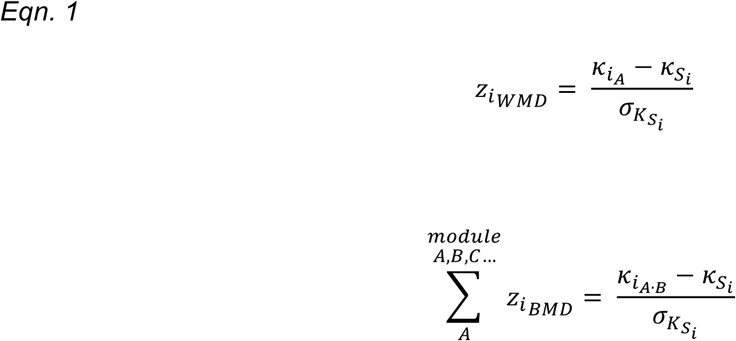

where 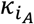 is the sum of all connectivity values for an ROI *i* within a module *A*, 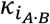 is the sum of all connectivity values for an ROI *i* between a module *A* and *B, 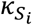* is the average of *κ* over all the nodes in *S*_*i*_, and 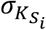 is the standard deviation of *κ*. in *S*_*i*_. This within-module degree z-score 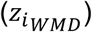 measures how well connected node *i* is to other nodes within the module, while the between-module degree z-score 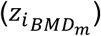 measures how well connected a node *i* within module *A* is to other nodes with another module (*B*, *C*, *ect*.) in the cortical parcellation. Here we sum the BMD score across all possible module pairings *A* · *B*, *A* · *C* etc., and then investigate the specific module-to-module BMD values that drive this mean, examining only nodes within the left PFC module (i.e., the stimulated module). As such, the calculation of the BMD is very similar to the participation coefficient, with the added benefit that BMD scores reflect the connections between *specific pairs* of modules, while the participation coefficient reflects the connectivity between a node and *all* modules outside of its local community. Thus, critical information is lost in the participation coefficient concerning the specific target of a node’s extra-module connectivity.

Nevertheless, to connect our novel measure to existing metrics, we include PC for all regions showing significant effects of memory success on BMD (see Table 1). As such, the calculation of 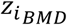 may be repeated for each module. In typical applications, nodes with a high *z*_*WMD*_ are interpreted to represent local, intramodular information-processing hubs, whereas nodes with high *z*_*BMD*_ show a relatively even distribution of connectivity across all modules. The central analysis focuses therefore on 4 functional connectivity adjacency matrices produced by this analytical pipeline: matrices for subsequently remembered or forgotten trials, after either 1Hz or 5Hz rTMS. As noted above, our statistical testing was based on examining differences between subsequently remembered and subsequently forgotten trials; given the number of statistical tests (411 cortical and subcortical ROIs), all comparisons were FDR-corrected using a Benjamini-Hochberg correction for multiple comparisons (Benjamini and Hochberg, 1995). We list these p-values in ascending order and denote them by *p*_(*k*)_, where *k* denotes the rank of a p-value. For a given *α* level, we find the largest *k* such that 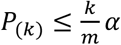, where *m* is the total number of hypotheses tested. Specifically, we applied this correction method within each modality (WMD, BMD), consistent with standard neuroimaging practice in which FDR corrections are applied within a specific contrast map, and not across multiple contrasts maps (Nichols et al., 2017).

**Table 1.**
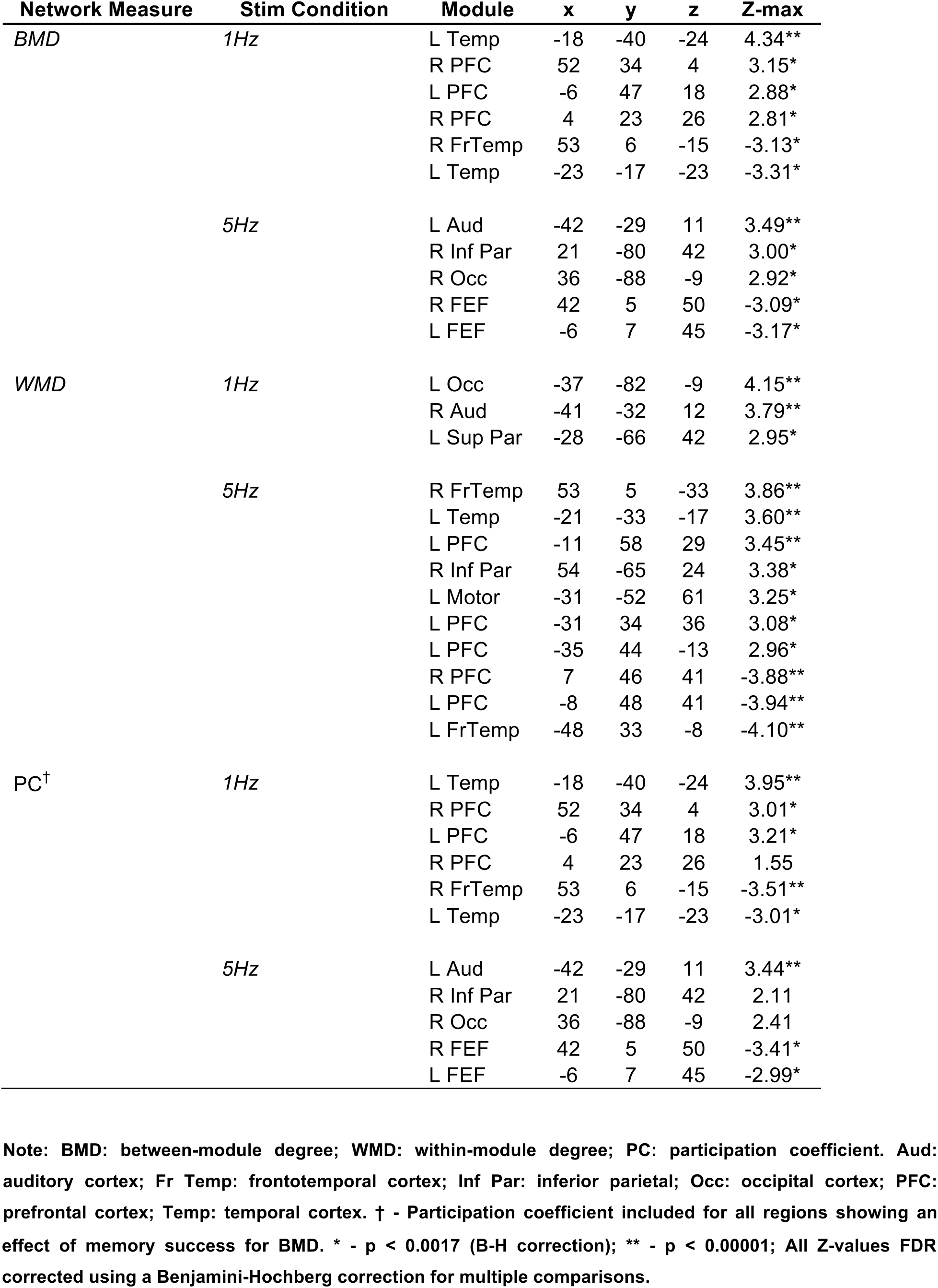
Regions demonstrating Subsequent Memory Effects on Network Metrics.

### Cross-modal comparisons

Lastly, we sought to test the hypothesis that the regions that are most affected by rTMS stimulation are best predicted by the structural connectivity with the stimulation site. A number of recent analyses have focused on first-order correlations between a structural connectivity matrix ***A_DTI_*** and a functional connectivity matrix ***A_fMRI_***, for a given region *R*, based on the assumption that direct structural connectivity should engender a corresponding modulation of functional connectivity (Zimmermann et al., 2016). The agreement of functional and structural connectivity between all regions was therefore calculated using a novel method of estimating the functional-structural relationship as a function of the structural path length in a structural matrix ***A_DTI_***. In order to enhance the effect of high connection weights (i.e., streamline counts) with respect to low values we use a nonlinear decreasing function *S* to map streamline count *C* to edge weight, *W*_*i,j*_ *= S(C*_*i,j*_) Following Everts et al. (2015), we use a decreasing sigmoidal function of the form

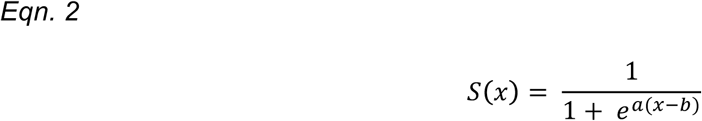

where *a* is a positive constant that determines the steepness of the sigmoid and *b* is a constant that determines the x-position of the steepest point of *S*.

We then calculated the Pearson’s product moment correlation between a vector describing the structural connectivity for both within-module connections 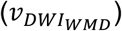 and between module connections 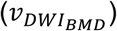, and vectors describing the corresponding vectors for functional connectivity *v*_*fMRI*_; given the role of negative correlations described above, we used the absolute value of the functional connectivity information for each condition. This cross-modal correlation was then repeated for both 1Hz and 5Hz functional matrices, using only the vector describing connectivity from the site of stimulation (as quantified by in-scanner fiducial markers). Statistical significance in both global and region-specific relationships were obtained by Fisher-transforming the *r*-values obtained for each subject and calculating paired-sample t-tests between 1Hz and 5Hz conditions; WMD and BMD were not compared directly, as these measures represent distinct distributions of connectivity values, with different numbers of contributing connections, and are thus not directly comparable.

## Results

### Behavioral performance

As explained in the Introduction, our goals were to investigate: (1) the effects of TMS on local activity and local/global connectivity, and (2) how these effects are influenced by structural connectivity. Consistent with our intended use of TMS as a physiological probe (Clapp et al., 2010; Feredoes et al., 2011), memory performance did not differ between rTMS conditions (mean d’ across subjects: 2.34 and 2.46 in the 1Hz and 5Hz rTMS conditions, respectively; t_13_ = 0.75, p = 0.6), nor did reaction times to correct trials (t_13_ = 1.19, p = 0.3). Furthermore, memory performance was not significantly different during either rTMS condition when compared to baseline memory performance (mean d’: 1.71), when compared either to 1Hz (t_13_ = 1.80, p = 0.11) or 5Hz rTMS conditions (t_13_ = 1.90, p = 0.08). This lack of a behavioral enhancement is critical, because it allowed us to infer that differences in network structure were uncontaminated by performance differences between 5Hz and 1Hz conditions.

### Effects of TMS on memory-related activity and connectivity

As expected, rTMS did significantly modulate memory-related activity and functional connectivity, as isolated by the subsequently remembered vs. forgotten contrast (subsequent memory effect—SME). Analyses on the effects of TMS and links to DTI measures focus on memory-related activity and connectivity (SMEs). Consistent with the compensation hypothesis, our first prediction was that *1Hz rTMS would reduce activity in the stimulated region (local activity) but increase its BMD (global connectivity), whereas 5Hz rTMS would increase both activity and WMD in the stimulated region.* We first investigated the effects of rTMS on activity in the stimulated brain region (left MFG) and then turned to rTMS effects on BMD and WMD.

#### rTMS effects on memory-related activity

Before focusing on the stimulated region, we performed a whole-brain analysis (see Figure 3). Compared to a baseline (no rTMS) condition, which primarily served as our functional localizer, 1Hz rTMS reduced SMEs in several brain regions, whereas 5Hz rTMS enhanced SMEs in multiple PFC, parietal, and ventral temporal areas typically associated with successful lexical encoding (Wagner et al., 1998; Prince et al., 2005). Consistent with our first prediction, the SME in the stimulated left MFG region (as shown in red in Figure 3B) was reduced by 1Hz rTMS compared to baseline (t_13_ = - 2.43, p < 0.01), but was increased by 5Hz rTMS (t_13_ = 2.83, p < 0.01) compared to baseline activation for subsequently remembered encoding trials. A pairwise comparison between 1Hz and 5Hz fMRI-estimated activity for successfully remembered trials was also significant (t_13_ = 3.43, p < 0.001). Having confirmed that rTMS modulated memory-related activity at our site of stimulation, we then turned to the effects of rTMS on BMD and WMD.

**Figure 3.**
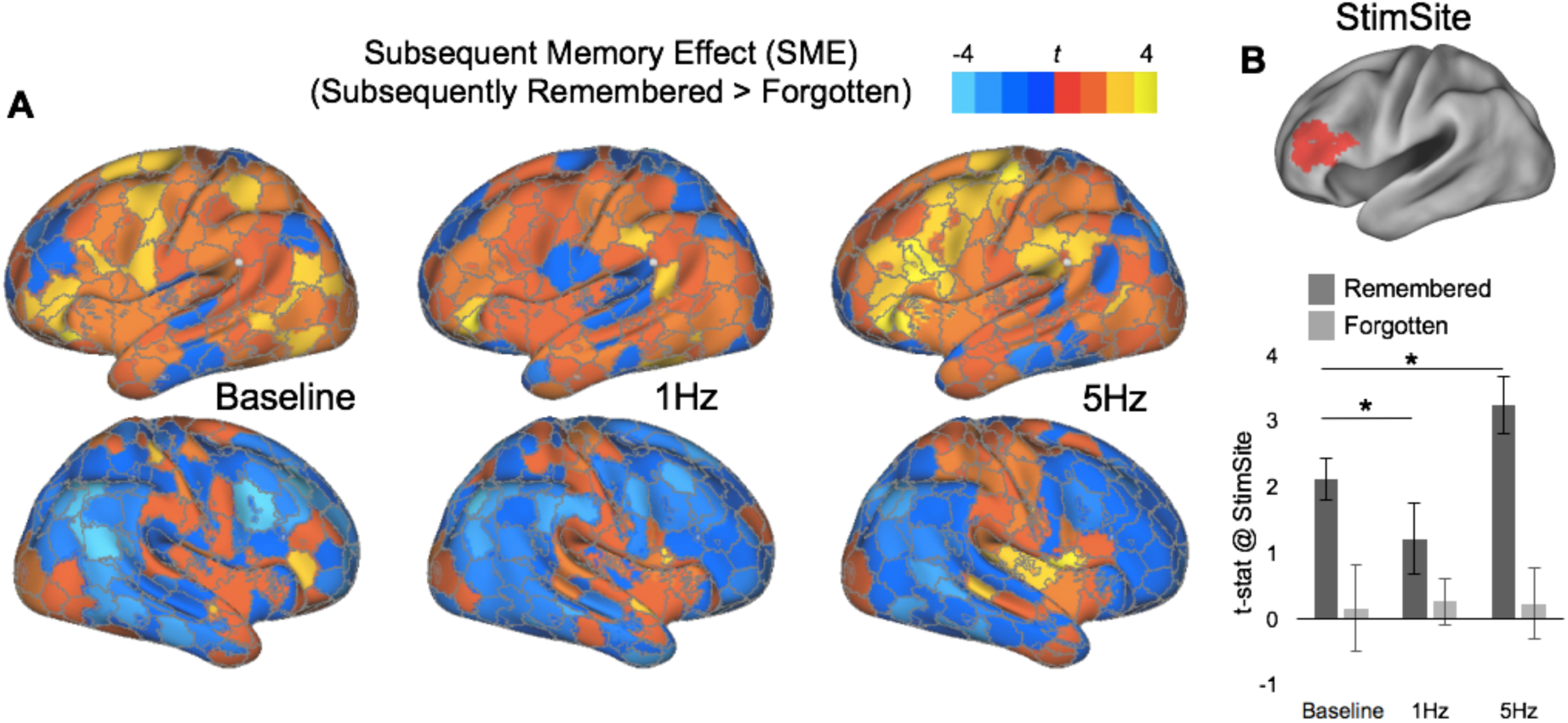
Effects of rTMS on subsequent memory effects. (SMEs = remember-forgotten). (A) Effects on 411 cortical ROIs during Localizer memory test. (B) The left PFC Stimulation Site within the left MFG is shown in red, with standard effects for subsequently Remembered and Forgotten trials during Baseline, 1Hz, and 5Hz conditions.

#### Multivariate Network Analyses

Our main analysis focused on the effects of excitation and inhibition the modular structure of memory success-related networks in the aging brain. We used 5Hz and 1Hz rTMS in order to enable a causal assessment of the stimulated brain region’s influence on connected brain regions, which we evaluate by summarizing the effects on the modular structure of success-related brain activity. We derived modularity for our sample based on a structural connectivity as assessed by DWI, and applied a data-driven algorithm for identifying consistent module partitions (Chang et al., 2012). The modularity partition based on diffusion tractography connectivity (adjusted streamlines) demonstrated strong modularity (Q = 0.71 across 100 permutations), with 16 distinct, symmetrical communities formed from the averaged structural connectome in our sample; community assignments are depicted in Figure 4. Each module comprises between 68 and 120 individual nodes. Within-module degree (WMD) quantifies functional connectivity among the nodes within each module, whereas between-module degree (BMD) assesses functional connectivity between different modules. We first considered the effects of rTMS on between-module degree (BMD).

**Figure 4.**
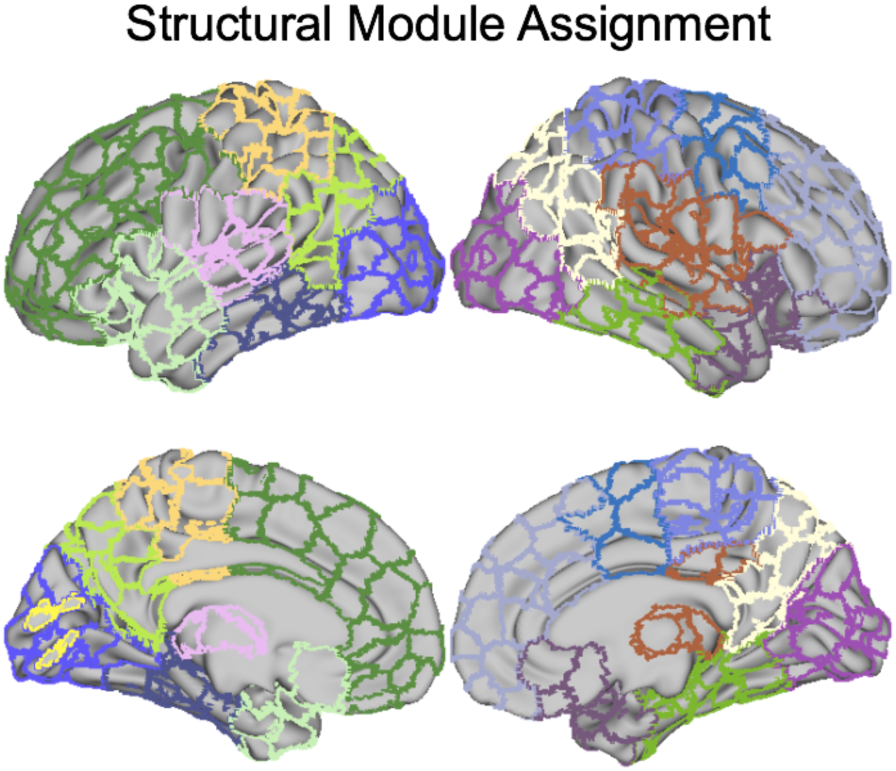
Structural module assignment. Border colors denote module assignment, and reflect module partitions based on structural connectivity patterns.

#### Effects of rTMS on between-module degree (BMD)

On the basis of the compensation hypothesis, we predicted that a 1Hz rTMS condition—which reduced local activity—would increase BMD both within local left PFC community, as well as with the more distant regions connected to these left PFC nodes. Consistent with this prediction, contrasts between successfully remembered and forgotten BMD in the 1Hz rTMS condition yielded large memory-related BMD *increases* in a number of bilateral PFC regions, left temporal cortex, as well as BMD *decreases* in bilateral fronto-temporal ROIs (Figure 5-middle; Table 1). In contrast with 1Hz rTMS, the same comparison in the 5Hz rTMS condition produced limited effects in auditory and occipital regions, as well as bilateral regions centered on the frontal eye fields (FEF; Figure 5-right).

**Figure 5.**
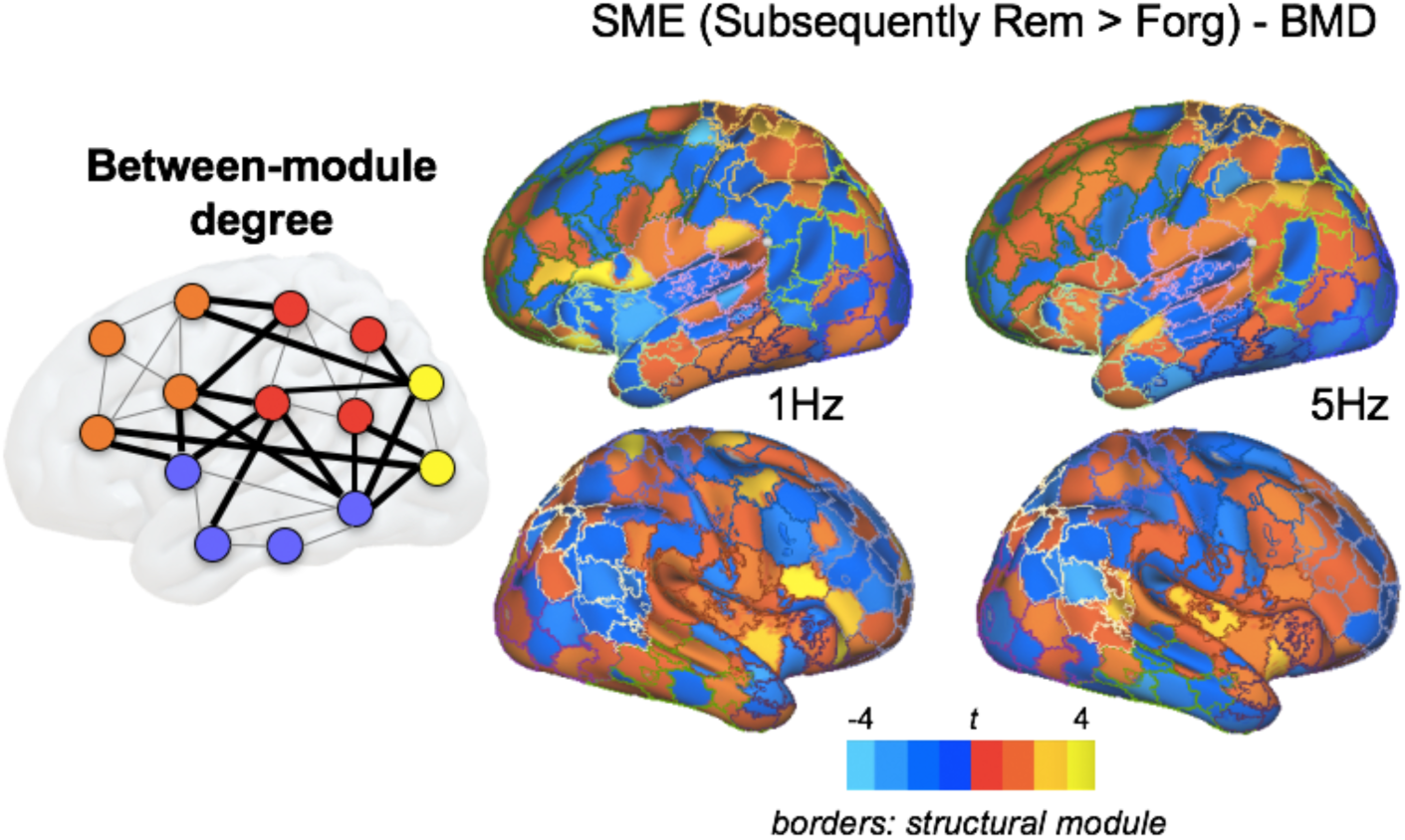
Between-Module Degree (BMD) associated with SME across rTMS conditions. use the same modularity defined by structural connectivity as above (A) in order to define effect sizes describing greater BMD for successfully remembered than forgotten trials. While connectivity after 5Hz stimulation demonstrates no PFC effect, BMD values in the 1Hz stimulation condition emerge in left and right PFC are greater for successful trials, suggesting that 1Hz stimulation is associated with more intermodular, global communication. Border colors denote module assignment.

To clarify the source of BMD increases in left PFC in the 1Hz rTMS condition, we calculated the proportion of BMD connections from left PFC to other modules (Figure 6). Although parietal regions make important contributions, the BMD increase in left PFC was primarily driven by its connections with right PFC, which accounted for 49% of all significant BMD connections. Modules comprising bilateral parietal cortex also contributed significantly to the left PFC BMD effect (Left Superior Parietal: 5%; Right Superior Parietal: 14%; Left Inferior Parietal: 18%; Right Inferior Parietal: 14%). Post-hoc tests showed that the BMD increases in left PFC were significantly due to enhanced connectivity with right MFG (mean BMD to right PFC module = 3.52; to all other modules < 1.0). It is worth noting that although right MFG was not directly stimulated with rTMS, it has strong white-matter connections with the stimulated left MFG ROI, which can explain the BMD effects. In sum, consistent with our first prediction, 1Hz rTMS on left MFG reduced its activity but enhanced its connectivity with the rest of the brain, particularly with contralateral right PFC regions. These findings provide direct support to the compensation hypothesis.

**Figure 6.**
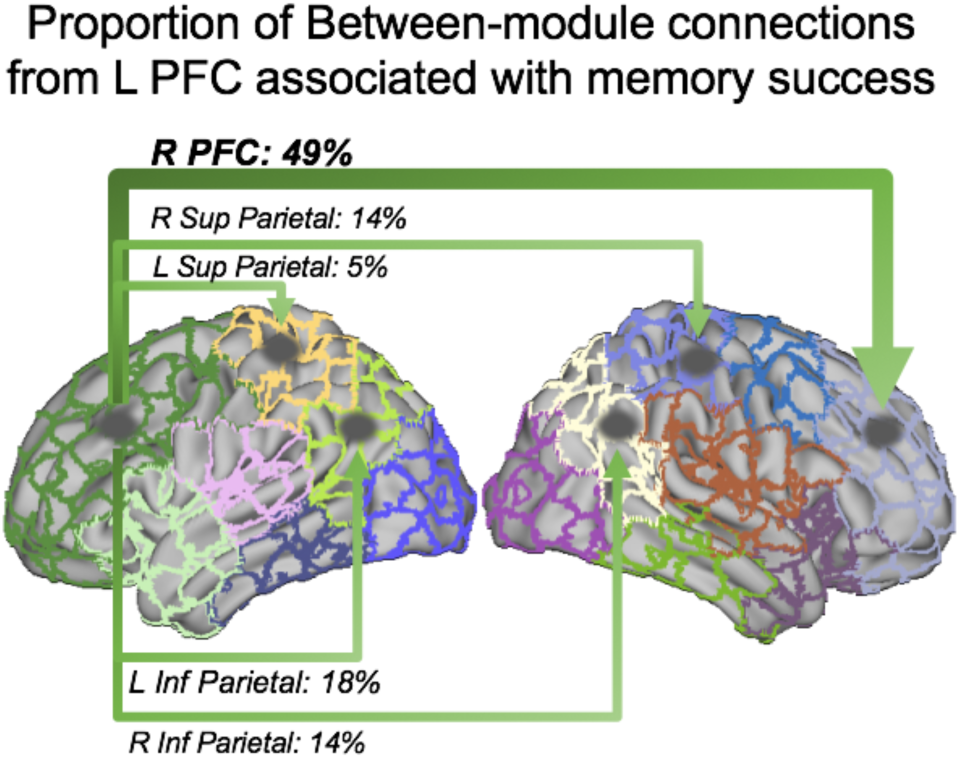
Proportion of BMD connections from Left PFC module to other modules. Between module connections in the 1Hz condition are largely attributable to a higher proportion of bilateral PFC connectivity to contralateral Right PFC module (49% of all between-module connections from the left PFC stimulation site), which also shows a success-related increase in BMD; See Figure 5.

Lastly, in order to explore the hypothesis that stimulation-induced changes in functional degree compensate for a deficit in core network regions, we examined both whole brain and stimulation site relationships between univariate SMEs (i.e., the increase in activity associated with successful encoding) and multivariate SMEs (i.e., the increase in between module connections (BMD) associated with successful encoding). Indeed, we found that 1Hz stimulation induced reliably negative relationships between univariate and multivariate measures, such that greater reductions in SMEs were associated with increases in the BMD. At the whole brain level, this relationship was much lower during the 1Hz than 5Hz (t_13_ = 9.75, p < 0.001) or baseline conditions (t_13_ = 10.64, p < 0.001; Figure 7A), and furthermore was more negative for successfully remembered than forgotten trials within the 1Hz condition (t_13_ = 10.02, p < 0.001; Figure 7B). At the level of the stimulation site, we also found a reliable negative correlation across subjects in subsequently remembered trials (*r*_*13*_ = −0.71, p < 0.05), which was absent during subsequently forgotten trials (*r*_*13*_ = −0.21, p < 0.05; Figure 7C). This effect was selective for the 1Hz rTMS condition, during which the local subsequent memory effect in fMRI univariate activity was attenuated (see Figure 3), and was not present after 5Hz rTMS, which had the opposite effect on local fMRI activity. No such relationships were observed in the relationship between univariate activity and WMD at the stimulation site (all *r* < 0.05), nor did whole-brain distributions show differences between stimulation conditions.

**Figure 7.**
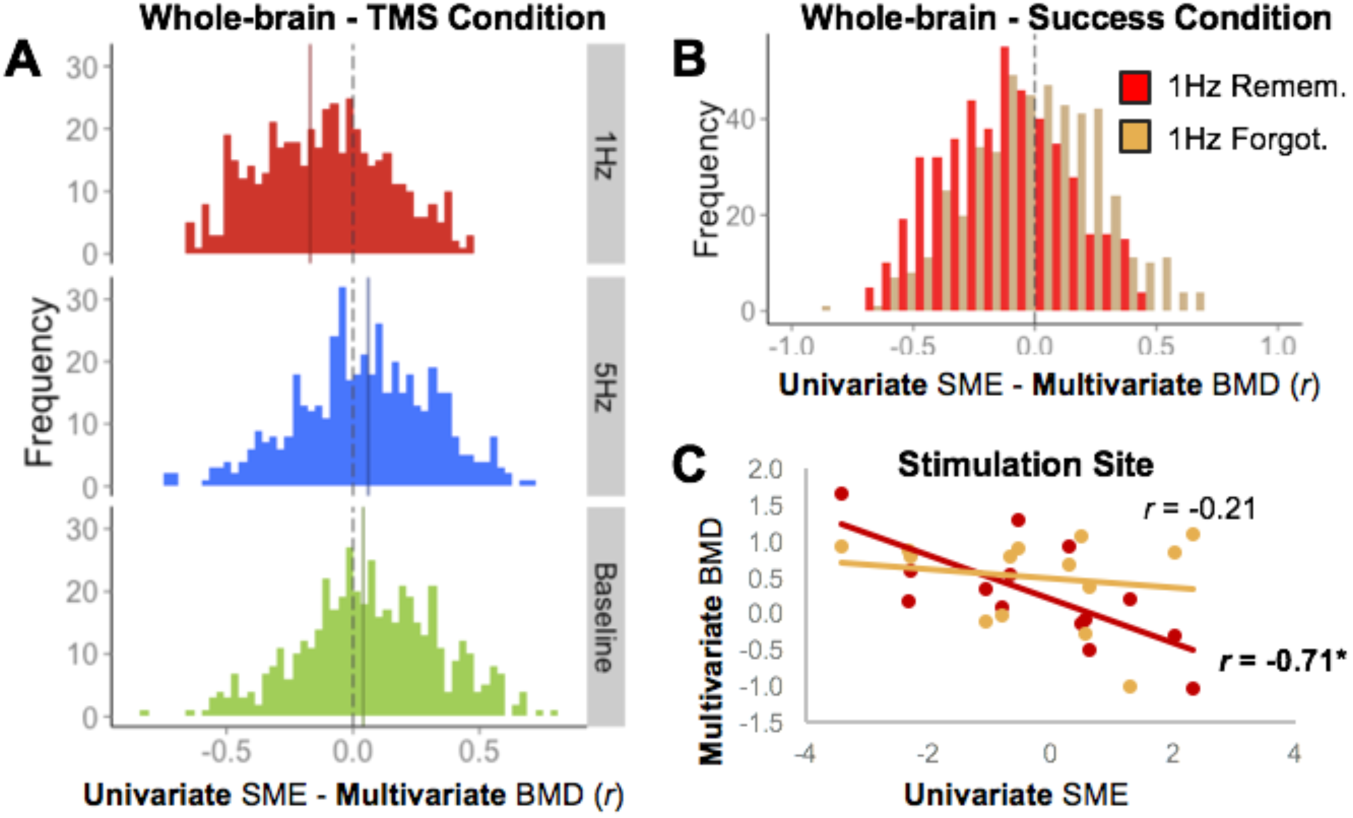
Relationship Between the Subsequent Memory Effect and multivariate Between-Module Degree. Distributions in (A) describe the correlation between SME and BMD, across subjects at each ROI. All distributions in this plot represent correctly remembered trials. The dashed line represents x = 0, while the solid like displays the mean of the distribution. (B) Distributions for subsequently remembered (red) and forgotten (tan) trials, demonstrating that this negative skew is related to successful memory encoding. (C) The effect is also present at the level of individual ROIs, in this case the site of stimulation. Overall this pattern of results suggests that as the SME is reduced by 1Hz rTMS, BMD-based connectivity increases to compensate for the local inhibition.

#### Effects of rTMS on within-module degree (WMD)

The second part of our first prediction is that a 5Hz rTMS condition that enhances activity in the stimulated region would lead to a WMD increase in this region. Consistent with this prediction, Figure 8 clearly shows that WMD in a number of ROIs near the stimulated left MFG region were enhanced by 5Hz rTMS, including the stimulation site (5Hz: t_13_ = 3.45, p < 0.005; 1Hz: t_13_ = 0.32, p = n.s.). Additionally, 5Hz rTMS also increased memory-related WMD in one right Frontotemporal ROI (t_13_ = 3.86, p < 0.001), an ROI in the Inferior Parietal module centered on the angular gyrus (t_13_ = 3.38, p < 0.005), and an ROI in the Left Temporal module centered on the left hippocampus (t_13_ = 3.60, p < 0.001), all regions typically associated with episodic memory success (Cabeza et al., 2008). Furthermore, 5Hz rTMS *decreased* memory-related WMD in one right PFC ROI (t_13_ = −3.88, p < 0.001) contralateral to the site of stimulation (x/y/z coordinates = 7/46/41), suggesting an isolation of the left hemisphere in the 5Hz condition. In contrast, 1Hz yielded only minor success-related WMD increases in auditory, occipital, and superior parietal modules (see Table 1 for a summary of all effects on BMD and WMD).

**Figure 8.**
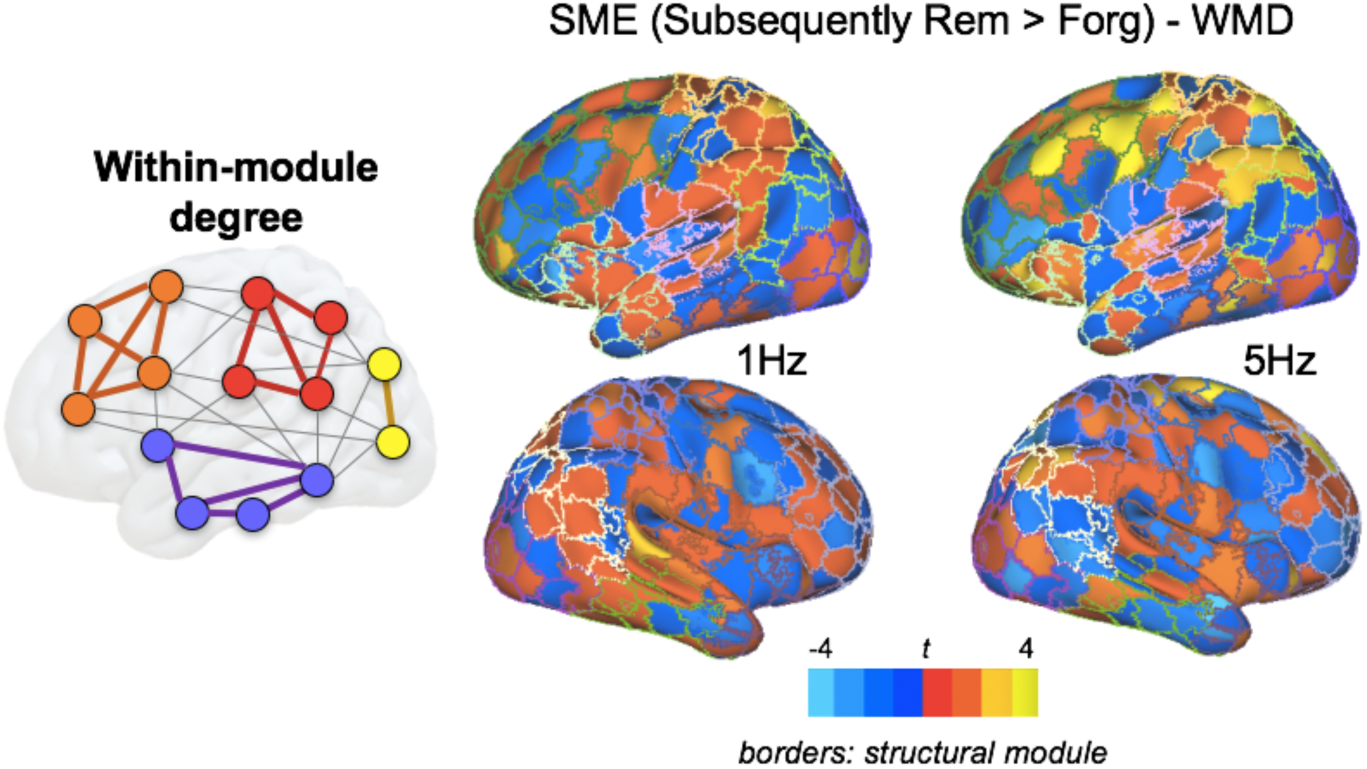
Within-Module Degree (WMD) associated with subsequent memory effects (SMEs) across rTMS conditions. Modularity defined by structural connectivity (Figure 4) describes a discrete network of left and right PFC modules, and premotor, motor, temporal, and occipital modules. An examination of the within-module degree for each ROI suggests greater local (left PFC) processing in the 5Hz condition. Border colors denote module assignment. In sum, consistent with the compensation hypothesis, we found that the 1Hz rTMS: (1) reduced the SME univariate activity in left MFG, and (2) increased WMD in left PFC, particularly between left PFC and contralateral right PFC regions. Lastly, we found that these two effects were connected to each other, as indicated by a negative correlation between the reduction in left MFG activity (local dysfunction) and the increase in WMD (global recruitment). The fact that this last effect occurred for remembered but not for forgotten trials is also consistent with the compensation hypothesis because it reflects the selectivity of this dynamic effect. In contrast, a 5Hz rTMS condition that enhanced local activity only led to an increase in local connectivity (WMD).

The overall focus of the present study was on examining the flexible dynamics associated with successful memory encoding using rTMS and fMRI. To ensure that the observed stimulation-induced network dynamics were specific to the encoding period, and not the subsequent retrieval period, we performed a post-hoc analysis in which we repeated our analysis of WMD and BMD measures during the episodic retrieval period following each encoding run. We found only one ROI within the left PFC module that demonstrated a significant main effect of memory success (Hits – Misses) in WMD (t_13_ = 2.87, p < 0.01; no significant BMD ROIs), and *no* significant main effect of Stimulation in either WMD or BMD that survived correction. This largely null finding can be attributable to the fact that rTMS effects in our study are transitory, and that the effect of 10 minutes of 5Hz or 1Hz rTMS is likely to dissipate within 20-25 minutes after the stimulation has terminated (Eisenegger et al., 2008).

### Cross-modal comparison

We found that distributed connectivity (BMD) of the stimulated left MFG region was enhanced by a 1Hz rTMS condition that reduced local activity, but not by a 5Hz rTMS condition that increased local activity. This finding is consistent with the idea that local dysfunction is counteracted by global recruitment (the compensation hypothesis). However, global recruitment depends on the strength of white-matter connections between distant brain regions. Thus, the effects of rTMS on functional connectivity should be predicted by the structural network along which stimulation-related responses might propagate, particularly in the 1Hz rTMS condition, which relies on a more distributed connectivity pattern. Thus, our second prediction was that *the global connectivity of the stimulated region should be correlated with structural connectivity in the 1Hz, but not the 5Hz, rTMS condition.*

To test this second prediction, we quantified the relationships between structural connectivity and the effects of 5Hz and 1Hz rTMS on memory-related functional connectivity for the stimulated left MFG region (x/y/z center of mass across subjects: −45/18/40, see Figure 9A). Pairwise t-tests revealed that for the within-module connections (red lines in Figure 9B), WMD at the stimulation site during 5Hz shows a modest correlation with structural connectivity to that site (r_13_ = .32), though this correlation is not different from the WMD-connectivity correlation in the 1Hz condition (r_13_ = .26). However, the increase in memory success-related WMD after 5Hz rTMS noted above suggests that the nature of the functional-structural association may be limited to the local structural patterns; a post-hoc test repeating the same correlations within a restricted set of left PFC regions within the left PFC module (see Figure 4) found similar results (WMD-Structural Connectivity @ 5Hz: r_13_ = .30; WMD-Structural Connectivity @ 1Hz: r_13_ = .29). For between-module connections, structural connectivity (i.e., the number of streamlines connecting two region pairs) was much more strongly correlated with the functional connectivity pattern for 1Hz than the functional connectivity pattern for 5Hz rTMS (t_13_ = 2.85, p < 0.01, yellow lines in Figure 9B). Outside left PFC, we did not find any significant difference between 1Hz and 5Hz rTMS on structure-function relationships. Thus, these effects describe an overall pattern consistent with our modularity results above, such that stimulation-related increases in local connectivity (i.e., WMD) after 5Hz rTMS, and increases in global connectivity (i.e., BMD) following 1Hz rTMS, were both constrained by short- and long-distance topology (respectively).

**Figure 9.**
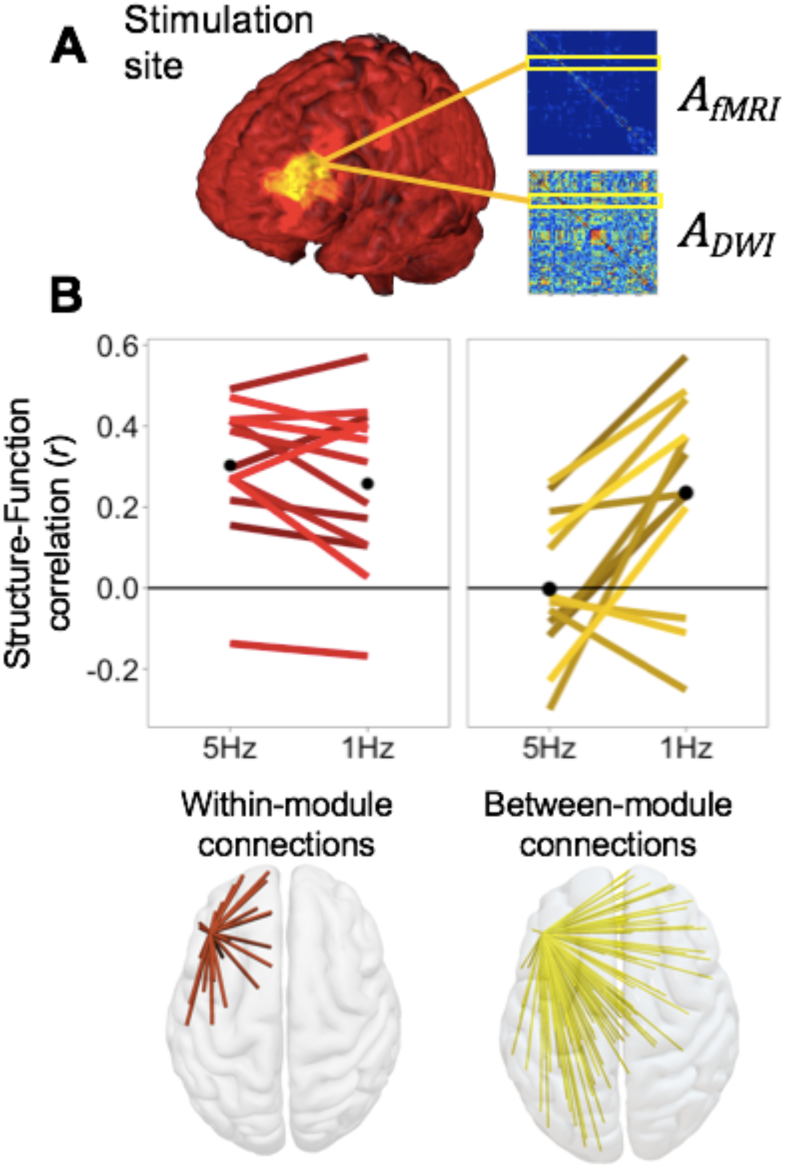
Structure-function relationships as a function of rTMS frequency and connection type. We represent the evolution of the correlation quality between structural and functional connectivity from a localized stimulation site (A) either *within* a local community, or *between* distinct modular communities. (B) Pairwise comparisons reveal consistently strong structure-function associations for network connections within the local module. In contrast, when considering between-module connections, this relationship was stronger for 1Hz than for 5Hz memory networks, suggesting the above increase in BMD connectivity associated with 1Hz rTMS is constrained by white matter connectivity. Note: pathways in B represent linkages as indexed by the graph theoretical measure of *distance*, and *not* direct tractography between regions.

## Discussion

Our overarching goal was to investigate the neural mechanisms linking local and global TMS effects by combining rTMS with fMRI and DTI. The study yielded two main findings. *First, a reduction in local activity was associated with an increase global connectivity.* Consistent with the compensation hypothesis, the 1Hz rTMS condition that caused under-activation at the stimulation site led to an increase in global connectivity (BMD). In contrast, the 5Hz rTMS condition that caused over-activation at the stimulation site yielded only an increase in local connectivity (WMD). *Second, the increase in global connectivity was modulated by structural connectivity.* In keeping with the idea that functional connectivity is constrained by white-matter health, global connectivity of the stimulated region was correlated with structural connectivity in the 1Hz, but not the 5Hz, rTMS condition. The sections below discuss these two findings and then consider their relations with aging.

### A reduction in local activity was associated with an increase in global connectivity

A critical finding from the current analysis is that 5Hz stimulation to a memory-specific target was associated with greater within-module connectivity (WMD) during successful encoding, while 1Hz stimulation engendered a more distributed pattern of connectivity with other modules (BMD). When examining the subject-wise relationship between univariate activity and BMD connectivity, we found a strong negative relationship for successfully remembered, but not forgotten, trials specific to the 1Hz condition, evidencing an adaptive relationship between local suppression of activity and more distant connectivity (Figs. 5, 6). This result suggests a highly responsive global network that is able to shift connectivity patterns in response to the disruption of local resources by relying on a more distributed pattern of connectivity (Vidal-Pineiro et al., 2014; Cocchi et al., 2015; Cocchi et al., 2016).

The dynamic relationship between local and distant connectivity is starting to be explored in a systematic way through the use of the graph-theoretic concept of “communities” or modules. A number of recent analyses have demonstrated that both weak intra-modular connections and more distant, inter-modular connections play a crucial role in establishing modular structure and predictive cognitive outcomes (Gallos et al., 2012; Santarnecchi et al., 2014). In these frameworks, as in our own, local interactions within a module characterize specialized functions (Sporns and Betzel, 2016), a trend that appears to develop through adolescence and across the lifespan (Power et al., 2010; Geerligs et al., 2015). Furthermore, the relationship between modularity and cognitive performance is typically positive, especially when the cognitive demands of the task can be localized to a discrete, processing system (e.g., working memory, as in Stevens et al., 2012) or when behaviors become more specialized over the lifespan (e.g., syntax, as in Meunier et al., 2014). Conversely, task operations which require greater global communication (and therefore reductions in functional modularity) are typically associated with cognitions requiring integration from multiple cortical communities, including visual awareness (Godwin et al., 2015) or episodic memory formation (Geib et al., 2015), which is consistent with global workspace models of brain function (Dehaene et al., 2011).

In the current study, we used static community boundaries to demonstrate how different stimulation conditions affect task-related memory processing within and across those boundaries. We showed that stimulation-induced reductions in local activity after 1Hz rTMS— but not 5Hz rTMS—resulted in increased connectivity between left PFC and other ipsilateral and contralateral modules in prefrontal and parietal regions (Fig. 7A), and that this effect was specific to successfully encoded trials (Fig. 7B, C), suggesting a frequency-specific role for this type of network reorganization. As noted above, frequency-specific TMS can selectively alter intrinsic neural dynamics between and within functionally specialized large-scale brain modules. This hypothesis is supported by the results of empirical and simulation studies, which suggest that the functional effects of focal changes in neural activity may extend outside a functionally segregated network (Bestmann et al., 2004; Alstott et al., 2009; van Dellen et al., 2013). Because large-scale modules are positioned at an intermediate scale between local and global integration, they may play a critical role in integrating local changes without reorganizing the backbone of large-scale brain communication.

### The increase in global connectivity was modulated by structural connectivity

Turning to our cross-modal network analysis, we found that the structural networks given by diffusion-weighted tractography reliably constrained the functional connectivity. We found that while within-module structural connectivity positively correlates with functional networks derived from both the post-5Hz and post-1Hz rTMS fMRI, the more long-range between-module connections more strongly predicted 1Hz than 5Hz functional success networks (Fig. 9). This pattern of results mirrors our earlier WMD/BMD findings, suggesting that global, between-module connectivity patterns are constrained by structural connectivity. While regional correlations between diffusion based structural connectivity and task-based functional connectivity remained consistently above chance—reflecting a general trend that structural connectivity may help explain some, but not all, of the variance in functional connectivity (Honey et al., 2009; Betzel et al., 2014)—this relationship was most pronounced in the connections emanating from the stimulation site. It is notable that the evaluation of BMD—a novel graph measure—provides a quantifiable measure of communication between specific bilateral PFC modules. The ability to successfully identify such a mechanism is therefore dependent on the explicit consideration to connectivity information between contralateral cortex (Davis and Cabeza, 2015). These measures may be lost in more conventional graph measures (e.g., PC), which make no distinction between specific long-range connections.

The finding that long-range connectivity (BMD) was selectively correlated with structural connectivity *in a specific stimulation condition* offers a mechanistic explanation for the anatomical basis for localizing the effects of changes at a proximal site at more distant locations in the aging brain. This mechanism operates on the intuitive principle that adaptive should prioritize cortico-cortical connections that are connected by white matter structure (Walhovd et al., 2011; Bartzokis et al., 2012). As such, we would expect stronger correlations between structural and functional connectivity in cross-hemispheric regions after inhibitory stimulation because they utilize available pathways (i.e., the genu of the corpus callosum) to contralateral cortical sites in the right PFC that facilitate memory functioning. This finding has important implications for the interpreting patterns of age-related change in functional activity, and suggests that adaptive reorganization in the aging brain takes advantage of existing structural architecture (Daselaar et al., 2014; Fjell et al., 2016). Thus, it seems now obvious that general, brain-wide conclusions about the relationship between structural and functional connectivity are limited, and any satisfying prediction about the adaptability of functional networks—in response to neurostimulation, injury, or age—must take into account the available network of structural connections to the site of insult. Therefore, there is sufficient evidence to suggest that these function-structure relationships are best characterized only by specific fiber systems during specific conditions. Furthermore, such multimodal relationships may not emerge when the functional networks are at rest, but only during active cognitive operations when those structural connections are necessary to the cognition at hand. In the context of our findings, such flexibility is in keeping with the expensive nature of long-range connections, as well as the dynamic nature of regional brain interactions.

Beyond increasing the ability to use combined brain imaging and stimulation to investigate brain function in normative populations, the present results suggest that DWI-based TMS targeting can extend the effectiveness of TMS in therapeutic applications (Luber et al., 2017). One obstacle for using TMS for effective treatments in memory disorders is that the brain region most critical for memory, the hippocampus, sits deep inside the brain, beyond the reach of direct TMS effects. The current finding suggests that cortical stimulation propagates to distant cortical and subcortical sites in a manner predicted by the 1^st^-order structural connectivity of the stimulation site. However, it remains unclear how these observed relationships may be affected by more large-scale system dynamics. Future work that focuses on the stability and changeability of network connectivity in memory states in the aging brain may therefore provide new information on the effectiveness of brain stimulation technologies as a therapeutic measure for cognitive decline.

### Both findings have implications for aging research

The finding that a reduction in local activity was associated with an increase in global connectivity is consistent with functional neuroimaging evidence that older adults who show deficits in specific brain regions (such as PFC), tend to display a more widespread pattern of brain activity than younger adults (Spaniol et al., 2009; Spreng et al., 2010). This effect is often attributed to compensatory mechanisms (Cabeza and Dennis, 2013; Reuter-Lorenz and Park, 2014), which could also explain the current rTMS findings. While typical assessments of these patterns are made across cohorts of younger and older adults, the current results represent a novel means of probing aging brain function by using frequency-specific changes in cognitive and network state. The impact of TMS on neural functioning in aging populations during memory tasks is still in its infancy. Nonetheless, new consensus is building towards the specific stimulation parameters that engender a positive effect on cognitive function in physiological and pathological aging (for review, see Hsu et al., 2015). A number of studies have elicited a positive role for TMS-induced excitability in older adult populations, largely utilizing excitatory 5-10Hz trains of rTMS (Luber et al., 2007; Eliasova et al., 2014; Brambilla et al., 2015) or theta-burst paradigms (Vidal-Pineiro et al., 2014).

Increased functional connectivity in healthy aged populations or in risk groups such as Alzheimer disease patients is often interpreted as a compensatory response to declining brain health (Agosta et al., 2012; Sheline and Raichle, 2013; Gomez-Ramirez et al., 2015), despite the fact that these increases are observed during resting state scans. Similarly, age-related examinations of graph-theoretic measures have typically relied on resting brain activity and generally describe a decline in the network cohesion (i.e., modularity) when compared with younger counterparts (Betzel et al., 2014; Cao et al., 2014). The utility of this task-free approach is limited. Growing evidence in task-related studies of whole-brain connectivity suggests that, despite this reduction in functional specificity, older adults are able to adapt the functional connectivity between functional networks (or modules) in order to adapt to task demands (Geerligs et al., 2014; Meunier et al., 2014). A recent analysis within- and between-network connections, using both resting and task-related activity, demonstrated that age-related increases in between-module connections were absent during rest (Grady et al., 2016), suggesting that task-related responses may be more sensitive to age-related changes in network functioning.

A critical component of characterizing a neural response as compensatory relies on the nature of its relationship to successful cognition. A growing number of studies have shown that noninvasive brain stimulation has a positive influence on various cognitive functions in both healthy and demented older adults (Hsu et al., 2015). These studies, which almost predominantly target left or right PFC suggest a positive role for the engagement of unilateral or bilateral (via spreading activation) PFC in maintaining healthy episodic memory function (Davis et al., 2012; Vidal-Pineiro et al., 2014; Brambilla et al., 2015). In line with these studies, we showed that stimulation-induced reductions in local activity after 1Hz rTMS resulted in increased connectivity between left and right PFC, suggesting a frequency-specific role for this type of network reorganization. As noted above, frequency-specific rTMS can selectively alter intrinsic neural dynamics between and within functionally specialized large-scale brain modules. This hypothesis is supported by the results of empirical and simulation studies, which suggest that the functional effects of focal changes in neural activity may extend outside a functionally segregated network (Bestmann et al., 2004; Alstott et al., 2009; van Dellen et al., 2013). Because large-scale modules are positioned at an intermediate scale between local and global integration, they may play a critical role in integrating local changes without reorganizing the backbone of large-scale brain communication. Nonetheless, the most effective parameters and dosing of stimulation (i.e., stimulation tool, frequency of rTMS, intensity, number of rTMS pulses, target brain region) in aging populations demands further investigation.

## Limitations

While the above findings provide a clear causal role for the role of bilateral brain dynamics in shaping healthy performance, the current analysis nonetheless suffers principally from two major limitations: a low sample size and a design that limits the interpretation of TMS-specific effects. With respect to the former limitation, the issue is whether the observed results might generalize to a broader aging population, or in fact is unique to older adults in the first place. It is important to note that although our hypothesis and two predictions mention OAs, they are not specific to aging. On the contrary, we believe the hypothesis that brain decline can be compensated with a shift from WMD to BMD would also apply with YAs suffering of brain dysfunction due to lesions or transient TMS effects. Thus, we could have investigated the hypothesis and predictions in either OAs or YAs. With respect to the latter limitation, we focused on elucidating a frequency-specific change in brain state (5Hz and 1Hz) in order to compare their effects on network organization in the aging brain, based on work from our own (Luber et al., 2007; Luber et al., 2008) and other laboratories (Thut et al., 2003; Peinemann et al., 2004; Schneider et al., 2010; Plow et al., 2014) which have found both fMRI-, EEG- and MEP-based evidence that 5Hz rTMS promotes increases in cortical excitability, while 1Hz has the opposite effect (though, see Caparelli et al., 2012). While our rTMS design lacked a same-day sham condition to eliminate the possibility that performance differences due to the arousing effects of TMS, neuroenhancement as such was not the goal of the present analysis. Our fMRI findings nonetheless support important distinctions between rTMS parameters, and suggest important frequency-specific effects on global network organization.

## Conclusions

The current analysis provides novel evidence that aging brains utilize a flexible set of neural dynamics to accomplish the same cognitive task under different stimulation conditions. Whereas 5Hz rTMS increased memory-related local connectivity (WMD), the application of 1Hz rTMS engendered more global connectivity (BMD) to different brain modules that were located bilaterally from the site of stimulation. These global effects were strongly constrained by structural connectivity derived from diffusion-weighted tractography. These results provide an integrated, causal explanation of the network interactions associated with successful memory encoding in older adults.

## Acknowledgements

This work was supported by National Institutes of Health Grant R01 AG19731.

